# Hypercholesterolemia Accelerates Both the Initiation and Progression of Angiotensin II-induced Abdominal Aortic Aneurysms

**DOI:** 10.1101/2020.01.03.893313

**Authors:** Jing Liu, Hisashi Sawada, Deborah A. Howatt, Jessica J. Moorleghen, Olga Vsevolozhskaya, Alan Daugherty, Hong S. Lu

## Abstract

**Objective:** This study determined whether hypercholesterolemia would contribute to both the initiation and progression of angiotensin (Ang)II-induced abdominal aortic aneurysms (AAAs) in mice.

**Methods and Results:** To determine whether hypercholesterolemia accelerates the initiation of AAAs, male low-density lipoprotein (LDL) receptor -/- mice were either fed one week of Western diet prior to starting AngII infusion or initiated Western diet one week after starting AngII infusion. During the first week of AngII infusion, mice fed normal diet had less luminal expansion of the suprarenal aorta compared to those initiated Western diet after the first week of AngII infusion. The two groups achieved comparable luminal dilation on week 2 through week 6 of AngII infusion as monitored by ultrasound. To determine whether hypercholesterolemia contributed to the progression of established AAAs, male LDL receptor -/- mice were fed Western diet and infused with AngII for 4 weeks. Mice with established AAAs were then stratified into two groups based on luminal diameters measured by ultrasound. While AngII infusion was continued for another 8 weeks in both groups, mice in one group were continuously fed Western diet, but diet in the other group was switched to normal laboratory diet. In the latter group, plasma cholesterol concentrations were reduced rapidly to approximately 500 mg/dl within one week after the diet was switched from Western diet to normal laboratory diet. Luminal expansion progressed constantly in mice continuously fed Western diet, whereas no continuous expansion was detected in mice that were switched to normal laboratory diet.

**Conclusions:** Hypercholesterolemia accelerates both the initiation of AAAs and progression of established AAAs in AngII-infused male LDL receptor -/- mice.

**Clinical Relevance:** Hypercholesterolemia is modestly associated with AAAs in observational or retrospective clinical studies. It is not feasible to study whether hypercholesterolemia contributes to the initiation of AAAs or progression of established AAAs in human. This study using AngII-induced AAA mouse model provides solid evidence that hypercholesterolemia contributes to both the initiation and progression of AAAs, supporting that statin therapy at any stage of AAA development may be beneficial to hypercholesterolemic patients with AAAs.

## INTRODUCTION

Hypercholesterolemia augments development of angiotensin II (AngII)-induced abdominal aortic aneurysms (AAAs) in male mice.^1-6^ This has been most commonly demonstrated in male low-density lipoprotein (LDL) receptor -/- mice fed a Western diet for one week prior to starting AngII infusion.^7^ This one week of Western diet feeding leads to profound increases of plasma cholesterol concentrations to more than 1,000 mg/dl.^3^ It is unclear whether a pre-existing hypercholesterolemic status is important to the initiation of AngII-induced AAAs, or it is critical for the progression of pre-existing AAAs.

Plasma total cholesterol concentrations are below 300 mg/dl in LDL receptor -/- mice fed normal laboratory diet, but increase to more than 1,000 mg/dl within a week of Western diet feeding.^3,8-10^ This LDL receptor -/- mouse in combination with diet manipulation provides a model to determine whether hypercholesterolemia contributes to the initiation and progression of AngII-induced AAAs. In this study using male LDL receptor -/- mice, we determined whether hypercholesterolemia would promote the initiation of AngII-induced AAAs, and whether maintaining hypercholesterolemia would modulate the progression of established AAAs.

## METHODS

The raw data that support the findings reported in this manuscript are available from the corresponding author upon reasonable request.

### Mice and Diet

Male low-density lipoprotein (LDL) receptor -/- mice (8 weeks old) were purchased from The Jackson Laboratory (Stock # 2207; Bar Harbor, ME, USA). Mice were housed in individually vented cages (5 mice/cage) on a light:dark cycle of 14:10 hours. The cage bedding was Teklad Sani-Chip bedding (Cat # 7090A; Envigo, Madison, WI, USA). Mice were fed a normal rodent laboratory diet (Diet # 2918; Envigo) and given drinking water from a reverse osmosis system ad libitum. To induce hypercholesterolemia, mice were fed a Western diet supplemented with saturated fat extracted from milk (21% wt/wt) and cholesterol (0.15% wt/wt supplemented and 0.05% wt/wt from the fat source; Diet # TD.88137; Envigo). Mice died of any reason prior to termination were excluded for data analyses.

In this study, we only used male mice because female LDL receptor -/- mice have very low incidence of AngII-induced AAAs.^3,11,12^ All mouse experiments reported in this manuscript were performed with the approval of the University of Kentucky Institutional Animal Care and Use Committee (University of Kentucky IACUC protocol number: 2006-0009).

### Mini Osmotic Pump Implantation and Angiotensin II Infusion

To induce AAAs, mice were infused with 1,000 ng/kg/min of AngII (Cat # H-1705; Bachem, Torrance, CA, USA) subcutaneously via Alzet mini osmotic pumps (Alzet Model # 2006; Durect Corp, Cupertino, CA, USA).^7^ In first study (Figure 1A), one group of mice started Western diet one week prior to AngII infusion (WD group), whereas the other group started this Western diet one week after initiating AngII infusion (ND - WD group). AngII infusion duration was 6 weeks for both groups.

**Figure 1.**
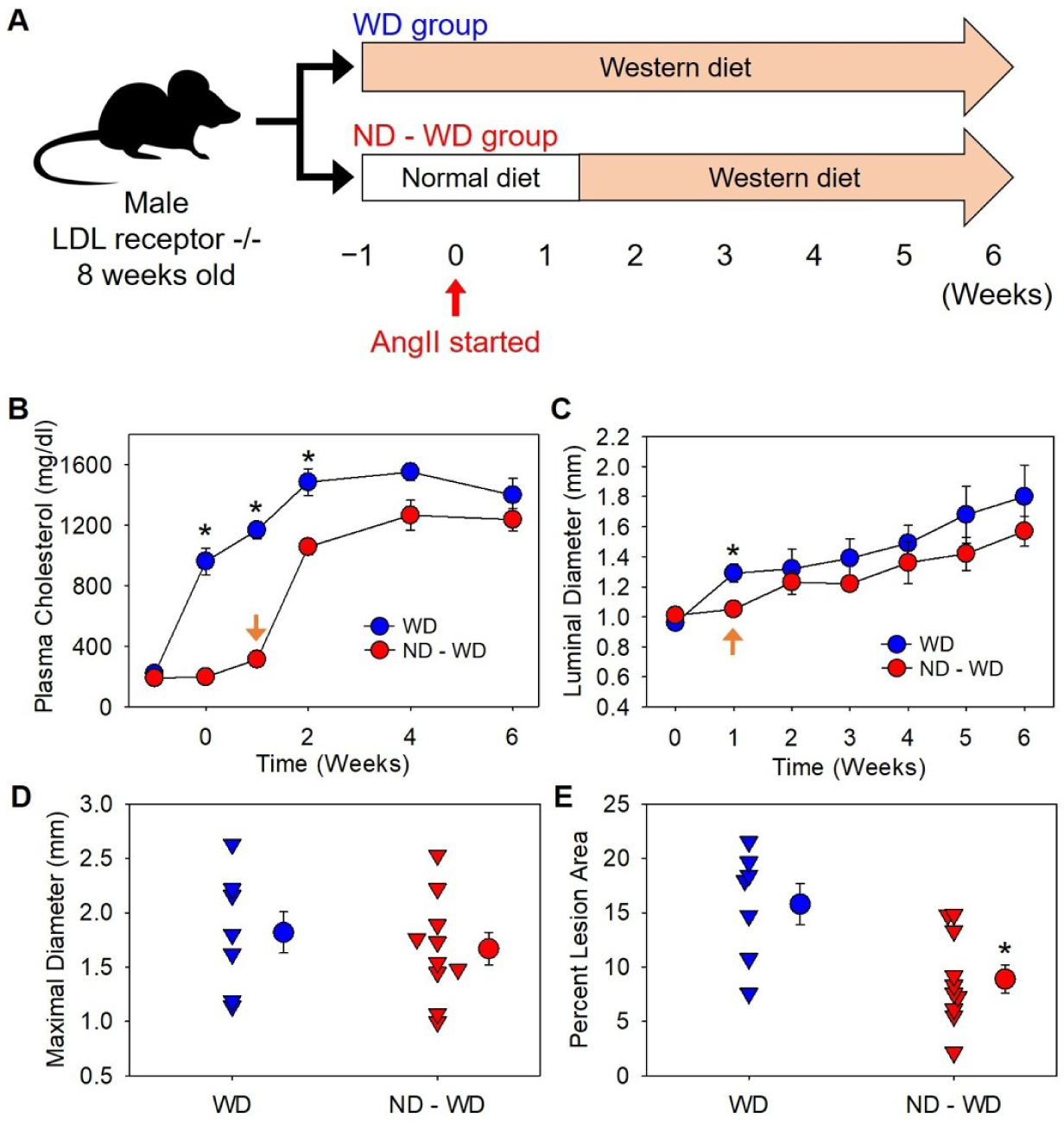
Hypercholesterolemia accelerated the initiation of AngII-induced AAAs in male LDL receptor -/- mice. (**A**) Mice in WD group initiated Western diet feeding one week prior to AngII infusion, and mice in ND - WD group initiated Western diet one week after starting AngII infusion. The duration of AngII infusion was 6 weeks. (**B**) Plasma cholesterol concentrations were measured using an enzymatic method. * P < 0.001, = 0.007, and < 0.001 between the two groups on Week 0, 1, and 2, respectively. (**C**) Maximal luminal diameters of suprarenal aortas were measured using ultrasound.* P = 0.004 on week 1. The orange arrow in (**B**) and (**C**) indicates the start of Western diet feeding in ND - WD group. (**D**) Maximal outer diameters of suprarenal aortas were measured using an ex vivo method. (**E**) Atherosclerosis was measured by an en face method. * P = 0.008. Triangles are values from individual mice. Circles represent means and error bars represent SEM.

In second study (Figure 2A), all mice were fed Western diet for one week prior to AngII infusion. Lumen diameters of suprarenal aortas were measured using noninvasive high frequency ultrasound system (Vevo 2100 with MS550D; FUJIFILM VisualSonics, Toronto, ON, Canada) at baseline and on day 28 of AngII infusion. AAAs were defined as 50% or more increase of the maximal lumen diameter of the suprarenal aorta on day 28 compared to the baseline (day 0). Based on luminal diameter measurements, mice exhibiting AAAs were stratified into two groups, and then either continuously fed Western diet (WD group) or switched to normal laboratory diet (WD - ND group). These two groups were infused with AngII for an additional 8 weeks. The entire duration of AngII infusion was 12 weeks achieved by implanting two mini osmotic pumps separately (Alzet Model # 2006): One pump was implanted on day 0 and replaced by a second pump on day 43. The second pump was preincubated in saline at 37 °C for more than 60 hours to permit immediate delivery of AngII after implantation of pumps in mice.

**Figure 2.**
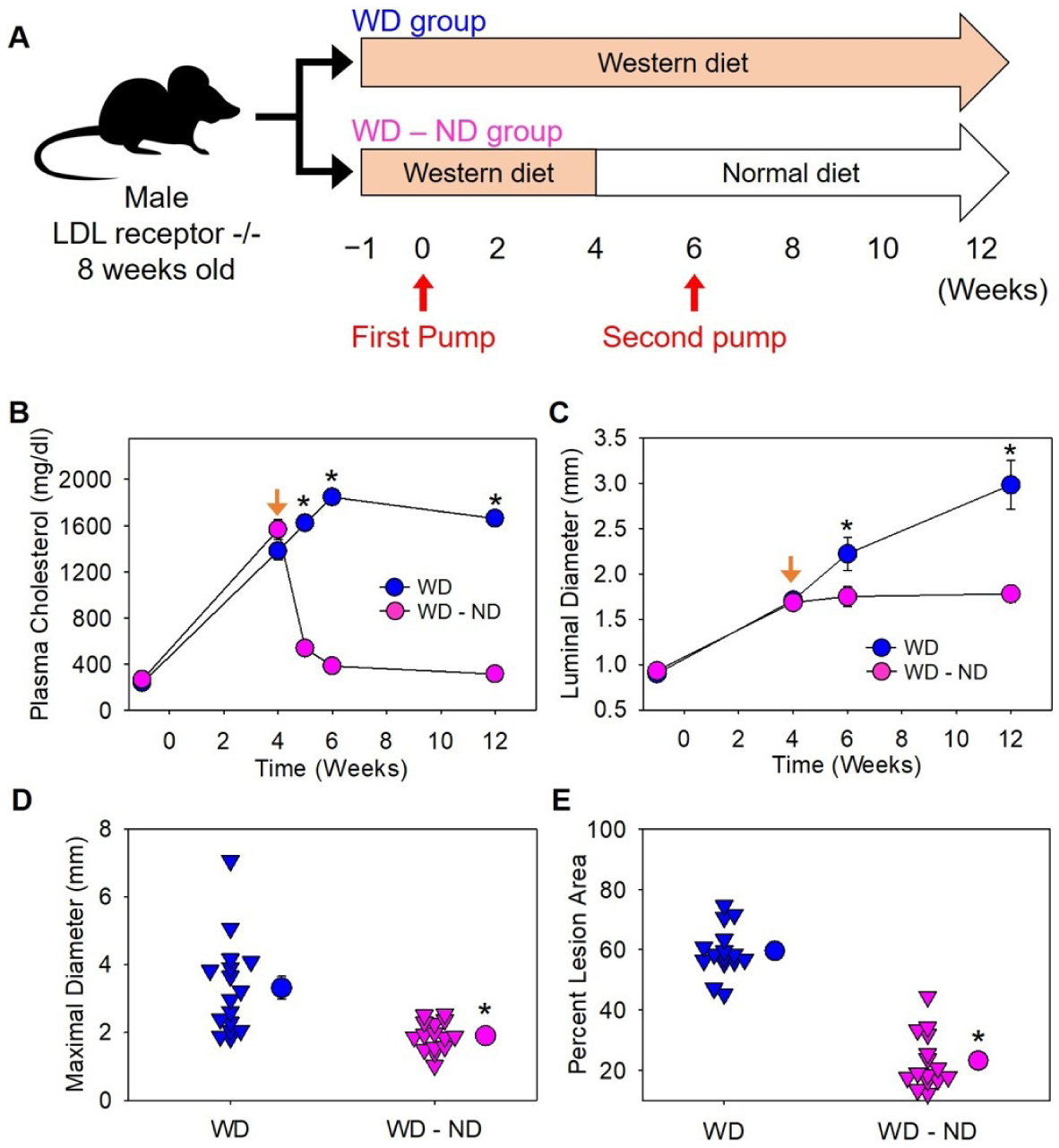
Hypercholesterolemia contributed to the progression of established AAAs in male LDL receptor -/- mice. **(A)** Mice in both groups started Western diet feeding one week prior to AngII infusion. Mice in WD group had been fed Western diet during AngII infusion, and diet for mice in WD - ND group was changed to normal laboratory diet after 4 weeks of AngII infusion. AngII infusion duration for both groups was 12 weeks. (**B**) Plasma cholesterol concentrations were measured using an enzymatic method. * P < 0.001 on week 5, 6, ans 12, respectively. (**C**) Maximal luminal diameters of suprarenal aortas were measured with ultrasound. * P = 0.02 and <0.001 on week 6 and 12, respectively. The orange arrow in (**B**) and (**C**) indicates diet was switched from Western diet to normal laboratory diet in WD - ND group. (**D**) Maximal outer diameters of suprarenal aortas were measured using an ex vivo method. * P < 0.001. (**E**) Atherosclerosis was measured by an en face method. * P < 0.001. Triangles are values from individual mice. Circles represent means and error bars represent SEM.

### Ultrasonography

Luminal diameters of suprarenal aortas were measured using a Vevo 2100 ultrasound imaging system at indicated intervals.^13^ Mice were anesthetized with isoflurane and restrained in a supine position. Short-axis scan was performed from the level of left renal artery moving vertically up to the suprarenal region. One hundred frames of cine loops were acquired and the maximal luminal diameter of the suprarenal aortic region was measured on images during aortic dilation phase. Luminal diameters were measured by one investigator, and verified by another investigator independently who were blind to study groups.

### Plasma Cholesterol Measurements

During each study, mice were bled consciously with submandibular bleeding. At termination mice were anesthetized using ketamine/xylazine cocktail, and blood samples were harvested by right ventricular puncture. All blood samples were collected with EDTA (final concentration: 1.8 mg/ml) and centrifuged at 400 g for 20 minutes, 4 °C to prepare plasma. Plasma cholesterol concentrations were measured using an enzymatic kit (Cat # 439-17501; Wako Chemicals USA, Richmond, VA, USA).

### Quantification of Aortic Dilation and Atherosclerosis

Aortas were dissected, fixed, cleaned, and pinned. Maximal outer diameter of the suprarenal aorta was measured ex vivo as a parameter for aortic dilation using Image-Pro software (Version 7; Media Cybernetics, Bethesda, MD, USA).

Thoracic aortas were cut open and pinned for quantification of intimal area and atherosclerotic lesion area in the ascending aorta, aortic arch and the proximal descending aorta using an en face technique.^14,15^ Atherosclerotic lesions, calculated as percent lesion area (lesion area/intimal area x 100%), were compared between groups following the recommended approach described in the AHA statement.^16^

### Statistical Analyses

Data are represented as means ± standard errors of means (SEM). To compare two groups on a continuous variable after termination, unpaired two-sided Student’s t test was performed for normally distributed and equally variant values, and Mann-Whitney rank sum test was used for variables not passing normality or equal variance test. Plasma cholesterol concentrations and luminal diameters of suprarenal aortas measured by ultrasonography at a series of time points were analyzed using linear mixed-effects models to compare trends over time among the sub-groups within the cohort, with random effects accounting for within-animal correlation. In general, unstructured correlation matrix with unequal variances was assumed to account for dependence among observations over time and within mice. Models were built using R version 3.3.2 statistical software. P < 0.05 was considered statistically significant.

## RESULTS

### Hypercholesterolemia accelerated the initiation of AngII-induced AAAs

Male LDL receptor -/- mice (N = 10/group) were randomized into two groups: WD group initiated Western diet one week prior to AngII infusion, and ND - WD group initiated this diet one week after AngII infusion was started (Figure 1A). Three mice in WD group were excluded for data analysis due to aortic rupture. Plasma cholesterol concentrations increased rapidly to approximately 1,000 mg/dl prior to the start of AngII infusion in WD group, whereas plasma cholesterol concentrations remained significantly lower within the first two weeks of AngII infusion in ND - WD group (Figure 1B). During the first week of AngII infusion, maximal luminal diameters of suprarenal aortas were smaller in ND - WD group than in WD Group as determined by ultrasonography (P = 0.0004). Luminal diameters became comparable between the two groups on 2 - 6 weeks of AngII infusion (Figure 1C), as were also validated by ex vivo measurements after termination (Figure 1D). Consistent with the less duration of hypercholesterolemia, atherosclerotic lesions were smaller in ND - WD group than in WD group (Figure 1E).

### Hypercholesterolemia augmented the progression of established AngII-induced AAAs

Seventy-nine male LDL receptor -/- mice were fed Western diet for one week prior to AngII infusion. The interval of AngII infusion was 4 weeks when Western diet continued. Eight mice died of aortic rupture during the 4 weeks of AngII infusion. Four weeks after AngII infusion 33 mice of the remaining 71 mice were stratified into two groups based on maximal luminal diameters (≥ 50% than baseline) of suprarenal aortas measured by ultrasound. Mice in WD group were fed Western diet continuously and diet for WD - ND group was changed from Western diet to normal laboratory diet (Figure 2A). AngII infusion continued for another 8 weeks among these 33 mice. Two mice were excluded for data analysis due to abdominal aortic rupture. Therefore, 16 mice in WD group and 15 mice in WD - ND group were included for data analyses. Plasma cholesterol concentrations decreased rapidly in WD - ND group within one week after diet change and remained low during the remaining interval of AngII infusion (Figure 2B). Maximal luminal diameters of suprarenal aortas continuously increased in WD group, but remained constant during the rest 8 weeks of AngII infusion in WD - ND group (Figure 2C). This significant effect of aortic diameters detected using ultrasound was confirmed after termination by ex vivo measurements (Figure 2D). Consistent with attenuation of hypercholesterolemia, atherosclerotic lesion sizes were smaller in WD - ND group than in WD group (Figure 2E).

## DISCUSSION

This study has demonstrated that hypercholesterolemia accelerates the initiation of AAAs and augments the progression of established AAAs in AngII-infused mice. This was demonstrated using male LDL receptor -/- mice fed normal laboratory diet or Western diet that rapidly change plasma cholesterol concentrations.^3^ This study not only confirmed previous reports that Western diet feeding quickly increased plasma cholesterol concentrations,^3,8,10^ but also provided evidence that plasma cholesterol concentrations reduced promptly after Western diet was withdrawn in LDL receptor -/- mice.

In this study, we used atherosclerosis as a “positive” control when we compared abdominal aortic dilation between groups because atherosclerosis has positive association with the magnitude and duration of hypercholesterolemia.^3,16,17^ Mice with either delayed Western diet feeding or removal of Western diet had less atherosclerotic lesions, which were consistent with lower plasma cholesterol concentrations or shorter period of hypercholesterolemia. In contrast, the magnitude of hypercholesterolemia did not affect development of AngII-induced AAAs, but removal of hypercholesterolemia attenuated the progression of established AAAs. These results support our previous findings that hypercholesterolemia augmenting AngII-induced AAAs has a threshold effect independent of the absolute concentration of plasma total cholesterol.^3^ In addition, the present study provided evidence that hypercholesterolemia accelerates AngII-induced AAAs during both the initiative and the progressive stages. These findings support that inhibition of hypercholesterolemia is critical to prevent AAA development and attenuate the progression of established AAAs.

Many studies used apolipoprotein E deficient mice to explore mechanisms of AngII-induced AAAs (a few examples from a large number of publications^2,18-22^). This hypercholesterolemic mouse strain is modestly hypercholesterolemic and does not need Western diet to accelerate AngII-induced AAAs.^3^ It appears that this mouse strain leads to higher incidence and mortality of AAAs, compared to LDL receptor -/- mice, although plasma cholesterol concentrations in apolipoprotein E -/- mice fed normal laboratory diet are much lower (∼ 300 - 400 mg/dl) compared to LDL receptor -/- mice fed Western diet (> 1000 mg/dl),^3^ implicating more complex mechanisms of AngII-induced AAAs in apolipoprotein E -/- mice. Therefore, LDL receptor -/- mice have benefits to study contributions of hypercholesterolemia and its related mechanisms to AngII-induced AAAs.

The present study provides guide for using AngII-induced AAA model: A pre-existing hypercholesterolemic condition accelerates AngII-induced AAAs in LDL receptor -/- mice. Therefore, our standard protocol to feed mice one week of Western diet prior to AngII infusion is optimal.^7^ Second, to continuously feed Western diet in LDL receptor -/- mice during AngII infusion is suggested since hypercholesterolemia is important for the progression of established AAAs. Dissecting effects of hypercholesterolemia in the human disease is a cost, effort, and time-consuming task, which involves many uncontrolled compound factors. Beyond being a guide how to use this AngII-induced AAA mouse model appropriately, this study, combined with our previous studies, also provides insights into understanding whether and how hypercholesterolemia contributes to AAAs in humans.^3,23,24^

### Conclusions

Hypercholesterolemia accelerates both the initiation and progression of AngII-induced AAAs.

## Author Contributions

Conception and design: AD, HSL

Analysis and interpretation: JL, HSL

Data collection: JL, DAH, JJM

Statistical analysis: HSL, OV

Raw data checking: HS

Writing the article: JL, HS, HSL

Editing the article: AD

Overall responsibility: HSL

All authors read the manuscript and approved the submission

